# Testing for Hardy-Weinberg Equilibrium in Structured Populations using NGS Data

**DOI:** 10.1101/468611

**Authors:** Jonas Meisner, Anders Albrechtsen

## Abstract

Testing for Hardy-Weinberg Equilibrium (HWE) is a common practice for quality control in genetic studies. Variable sites violating HWE may be identified as technical errors in the sequencing or genotyping process, or they may be of special evolutionary interest. Large-scale genetic studies based on next-generation sequencing (NGS) methods have become more prevalent as cost is decreasing but these methods are still associated with statistical uncertainty. The large-scale studies usually consist of samples from diverse ancestries that make the existence of some degree of population structure almost inevitable. Precautions are therefore needed when analyzing these datasets, as population structure causes deviations from HWE. Here we propose a method that takes population structure into account in the testing for HWE, such that other factors causing deviations from HWE can be detected. We show the effectiveness of our method in NGS data, as well as in genotype data, for both simulated and real datasets, where the use of genotype likelihoods enables us to model the uncertainty for low-depth sequencing data.

## 1 Introduction

Genotype frequencies in a population are normally described using the principle of Hardy-Weinberg Equilibrium (HWE) [1, 2]. Under the assumption of HWE, genotype frequencies can be defined as functions of allele frequencies which are conveniently captured as the Binomial distribution for diallelic sites. The HWE states that genotype and allele frequencies will remain constant in non-overlapping generations in the absence of other evolutionary forces given an assumption of random mating. Testing for HWE in a population has therefore become a very common tool for detecting possible sequencing or genotyping errors, population stratification as well as other effects leading to non-random mating, acting as a quality control step in genetic analyses [3, 4]. Extensions to HWE have been defined to incorporate an inbreeding coefficient in the statistical models to quantify deviations from HWE as a deficiency in observed heterozygotes. However, population structure will also lead to violations of the expected Hardy-Weinberg (HW) proportions by increasing the observed homozygosity due to the Wahlund effect, or increasing the observed heterozygosity due to recent admixture.

Recent methods have been developed to account for population structure using individual allele frequencies estimated from principal component analysis (PCA) [5–7]. PCA is a commonly used tool in population genetics for inferring population structure, as it has an advantage of describing individuals along axes of genetic variation instead of having to assign them in clusters [8]. The individual allele frequencies represent probabilities of the distribution from which the genotypes of each individual are sampled from given their inferred population structure [5]. These methods have shown to be very effective in large datasets with diverse ancestries, where population structure can be naturally taken into account using principal components. We have recently demonstrated the effectiveness of individual allele frequencies in next-generation sequencing (NGS) data as well, where we accurately infer population structure using an iterative algorithm in low-depth sequencing data [7].

NGS data has become more prevalent in genetic studies as the cost of whole-genome sequencing has decreased [9–12]. This has also lead to an increased amount of large-scale sequencing studies of samples with diverse ancestries [13, 14]. But sequencing depth is usually lowered to meet the demand of the large sample sizes in these large-scale projects, which comes at the price of introducing uncertainty into genotype calls. Probabilistic methods have therefore been developed to account for this uncertainty, and population genetic parameters are modelled using genotype likelihoods to retain information of the sequencing process. This has been shown to improve inferences for low- and medium-depth sequencing data (< 15X) [7, 15–18]. Likewise for genotype data, it is possible to model the missing genotypes instead of the standard practices of assuming no structure.

In this study, we propose a method to test for HWE in structured populations on the basis of genotype likelihoods. The method incorporates individual allele frequencies to account for population structure, such that we are able to test for effects leading to non-random mating other than population structure. Our method is implemented into the PCAngsd framework [7] for ease of use with both NGS and genotype data. We demonstrate its usefulness in both simulated and real datasets.

## 2 Materials and methods

It is assumed that individuals are diploid and variable sites are diallelic with genotypes coded as the number of minor alleles, *g* = {0, 1, 2}, and that the major and minor alleles are known for a dataset of *n* individuals and *m* sites. Based on individual allele frequencies, the genotypes of the individuals are assumed to be sampled as follow given their inferred population structure, under the assumption of HWE:

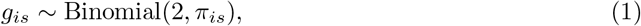

with *π_is_* being the individual allele frequency for individual *i* in site *s*. Individual allele frequencies were introduced in `STRUCTURE` [19] based on admixture proportions, population-specific allele frequencies and an assumption of *K* ancestral populations, however recent methods have employed approaches to estimate individual allele frequencies from the inferred population structure using PCA instead [7, 20].

However, the genotypes are not observed in sequencing data and we will work directly on genotype likelihoods instead. The genotype likelihood, *P*(*X_is_ | G* = *g*) can be described as the probability of observing the sequencing data *X_is_* given the genotype *g* for individual *i* in site *s*. We are therefore proposing a method for estimating per-site inbreeding coefficients and computing likelihood ratios to test for HWE. The method is an extension to an expectation-maximization (EM) algorithm derived in Vieria et al. (2013) [18] using genotype likelihoods. The EM algorithm is based on Wright’s coefficient of inbreeding for a site *s* defined as:

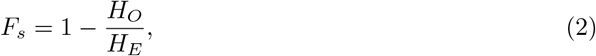

where *H*_0_ is the observed number of heterozygotes and *H_E_* is the expected number of heterozygotes. We extend the model by substituting population allele frequencies for individual allele frequencies in the likelihood function of the model to take population structure into account. In this way, we are able to estimate per-site inbreeding coefficients in structured and admixed populations. The likelihood of the inbreeding coefficient in a site *s* is defined as follows by assuming independence between individuals conditional on the population structure captured by individual allele frequencies:

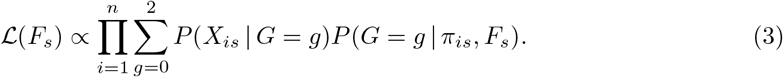

Here *F_s_* is the per-site inbreeding coefficient, *π_is_* is the individual allele frequency and *X_is_* is the observed sequencing data for an individual *i* in a site *s*. The genotype probability, *P*(*G* = *g | π_is_, F_s_*), is computed from Hardy-Weinberg proportions with the inbreeding coefficient incorporated to model deviations from HWE. Thus, for *G* = 0, 1, 2:

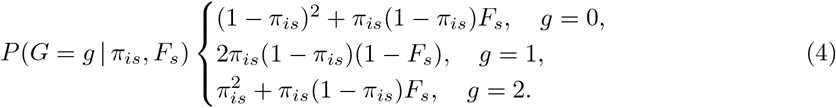

In this likelihood framework, *F_s_* is normally restricted to [0, 1], however we allow it to be in the interval of [−1, 1], where a negative estimate indicates an excess of heterozygosity and a positive estimate indicates an excess of homozygosity in site *s*. While a positive inbreeding coefficient does not change the allele frequency, a negative inbreeding coefficient will increase the sample allele frequency as the fraction of heterozygous individuals increases. A fixed allele frequency and a negative inbreeding coefficient can lead to negative probabilities for the homozygous genotypes in Equation 4 and thus make the distribution invalid. For example, *F_s_* = −1 can only have an allele frequency of 0.5 since all individuals will be heterozygous. In order to prevent this, we propose a heuristic distribution by truncating negative frequencies to 0 and rescaling the distribution to sum to one. The results of the truncation is visualized in Figure S1. *F_s_* has no biological interpretation as an inbreeding coefficient for negative values, but instead only act as some measure of deviation from HW proportions in the direction of excess of heterozygosity. It is therefore noteworthy that *F_s_* will not be behave the same in the negative domain as for the positive. We will still refer to *F_s_* as per-site inbreeding coefficient for convenience.

The likelihood is maximized using the proposed EM algorithm. The EM algorithm is fully described in the supplementary material. Using the maximum likelihood estimate, we construct a likelihood ratio test (LRT) statistic, *D_s_*, to test for deviations from HWE in each site. The null model is defined as *F_s_* = 0 and the alternative model is defined as the maximum likelihood estimate, 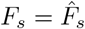:

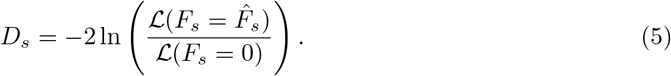

*D_s_* will be χ^2^ distributed with 1 degree of freedom.

### 2.1 Simulation of genotypes and sequencing data

We have simulated genotypes and low-depth sequencing data to test the capabilities of our method. Using allele frequencies of the reference panel of the Human Genome Diversity Project (HGDP) [21] from three populations (French, Han Chinese, Yoruba), we have simulated genotypes of 330 individuals using a binomial model. 100 individuals have been simulated from each of the 3 populations, while 30 individuals have been simulated with different degrees of admixture between pairs of the three populations based on varying admixture proportions to represent admixed individuals. The data consist of 340K variable sites and linkage disequilibrium has not been simulated between sites. Low-depth sequencing data and genotype likelihoods are simulated from the sampled genotypes based on a previously used approach [7, 17, 22]. For one individual, the number of reads in each site is sampled from a Poisson distribution with mean *d_i_*. Here *d_i_* represent the average sequencing depth of individual *i*, which is sampled as *d_i_ ∼ N*(5, 1). Thus, the sequencing depth of the simulated sequencing data is ∼5X. The simulated genotypes are then used to sample the number of reads being the minor allele based on a binomial model also using *d_i_* as parameter. Finally, the genotype likelihoods of the three genotypes are obtained from the probability mass function of the binomial distribution. Sequencing errors are incorporated using 0.01 as the probability of a sampled read being an error.

The data generated from the described procedure is regarded as Scenario 1, where all sites are sampled in HWE. However we also generate a second scenario, regarded as Scenario 2, where half of the sites deviate from HWE. This is done by changing the observed heterozygous genotypes to either of the homozygous genotypes with a probability of 0.5, such that *F* = 0.5 for a quarter of the sites. Likewise in the opposite direction, we change the homozygous genotypes to being heterozygous with a probability of 0.5 in order to simulate *F* = −0.5 for a different quarter of the sites. But the latter case will cause changes in allele frequencies due to imbalance between the numbers of homozygous genotypes, such that *F* = −0.5 does not when using sample allele frequencies.

### 2.2 1000 Genomes Project data

We also test our method on two genotype and sequencing data of the phase 3 release from the 1000 Genomes Project [13, 14]. The dataset consists of 366 unrelated individuals from four populations with 56 individuals from ASW (Americans of African Ancestry), 99 from CEU (Utah residents with Northern and Western European ancestry), 103 from CHB (Han Chinese in Beijing) and 108 from YRI (Yoruba in Ibadan). The genotype data is based on variant calls that consists of 7.4 million variable sites after data filtering, and the sequencing data is based on low coverage NGS data of the same individuals in 7.9 million variable sites with 6.9 million overlapping with the genotype data. The average sequencing depth in the low coverage dataset is estimated to 6.1X (varying from 1.7 − 13.6X) based on the used filters. Data filtering and generation of genotype likelihoods from the low coverage dataset have been performed in ANGSD [23]. The filtering options used for both datasets are described in the supplementary material.

### 2.3 Data accessibility

The method is integrated in the PCAngsd framework which is freely available at https://github.com/Rosemeis/pcangsd. The datasets used from the phase 3 release of the 1000 Genomes Project [13, 14] are available at ftp://ftp.1000genomes.ebi.ac.uk/vol1/ftp/ and a list of the used individuals is included as supplementary material.

## 3 Results

We have implemented our method into the PCAngsd framework and will also refer as to in the following results. Through the framework, the method will work on both genotype likelihoods in Beagle format as well as standard genotype data in binary PLINK format. PCAngsd converts the genotype matrix into the genotype likelihood format on-the-go for ease of use. The user can also incorporate an error model in the observed genotypes to factor in the possibility of alleles being errors. Here we apply our method to both genotype likelihoods and genotype data in simulated and real datasets using *ε* = 0 for comparison reasons.

Another method for testing for HWE in structured populations has recently been proposed by Hao and Storey in their software sHWE [24], which also uses individual allele frequencies, but only works on genotype data. However, our method has the advantage of estimating per-site inbreeding coefficients in addition to a test statistic as well as working on both sequencing and genotype data. We compare our method to both sHWE and the commonly used implementation in PLINK [25] which does not accommodate population structure.

### 3.1 Simulations

As proof of concept, we have applied our method to simulated genotype and low-depth sequencing data of 330 individuals from three different populations (French, Han Chinese, Yoruba) in 340K variable sites. The inferred population structure using PCAngsd is visualized in Figure 1 and we have used the top two eigenvectors to model the individual allele frequencies. As described in the *Materials and methods* section, two scenarios have been simulated. Scenario 1, where none of the sites are sampled to be out of HW proportions (*F* = 0), and Scenario 2, where half of the sites are sampled with *F* = 0 while the other half are equally sampled with either *F* = 0.5 or *F* = −0.5 being out of HW proportions.

**Figure 1:**
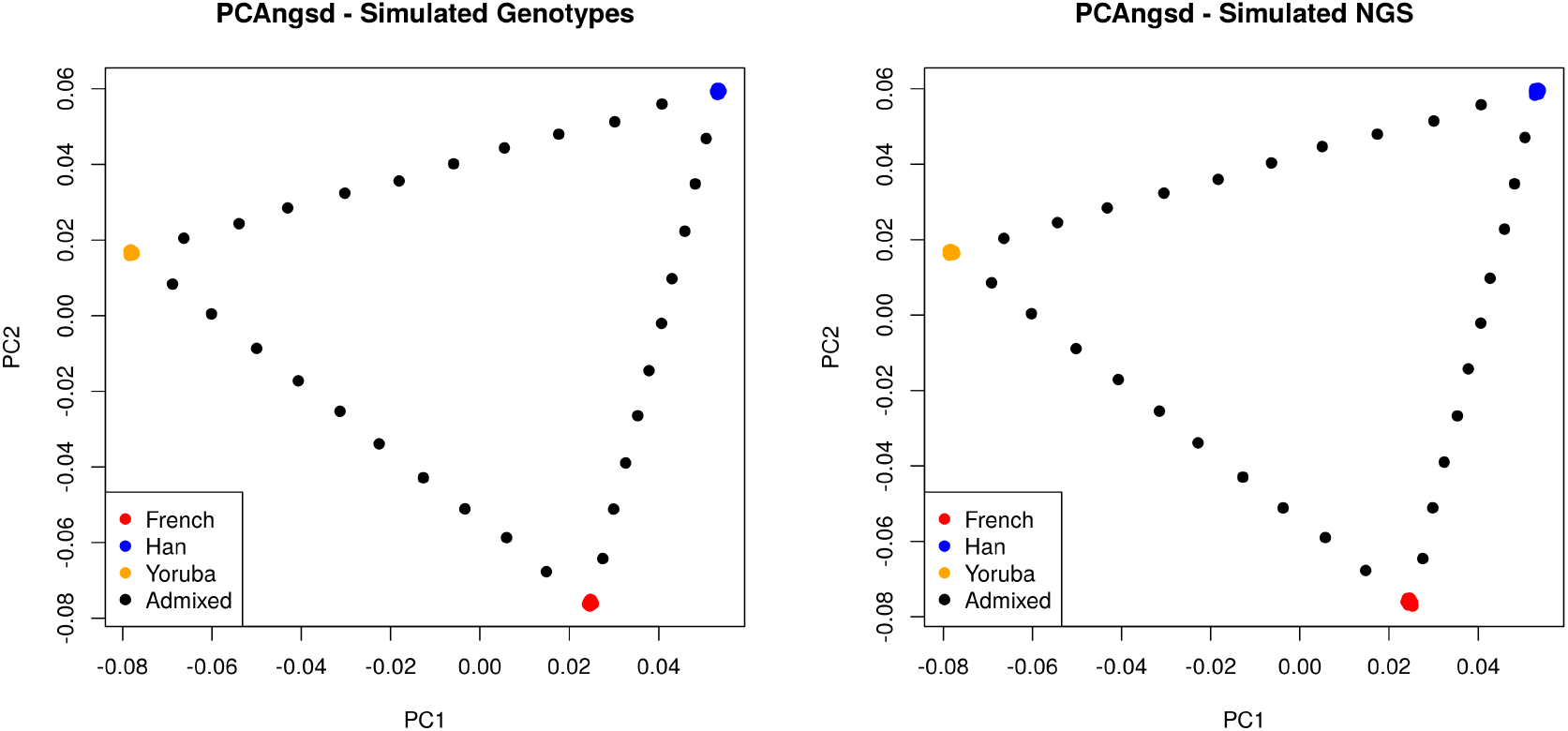
PCA plots of the simulated data. The inferred population structure of the left plot is generated from the simulated genotypes, and the right plot is generated from the simulated genotype likelihoods.

In Scenario 1, we are able to estimate per-site inbreeding coefficients that nicely follow a normal distribution around the expected value (*F* = 0) with the spread representing sampling variance, displayed in Figure 2. As seen by the QQ plots in Figure 3, our test statistic is also behaving as expected for both genotype and low-depth sequencing data. PCAngsd performs very similarly to sHWE, while the test statistics of PLINK are inflated and biased due to not taking population structure into account. sHWE is used with three logistic factors, as one factor represents the intercept. sHWE and PLINK are also applied to naively called genotypes of the sequencing data in Scenario 1 to demonstrate the difficulties in analyzing low-depth sequencing data. Genotypes are simply called by choosing the genotype with the highest likelihood, and the results are displayed in Figure 4. Here it is clearly seen that the two methods have inflated test statistics as the statistical uncertainty in the genotypes are not taken into account, where as PCAngsd is able to account for this uncertainty by working directly on genotype likelihoods.

**Figure 2:**
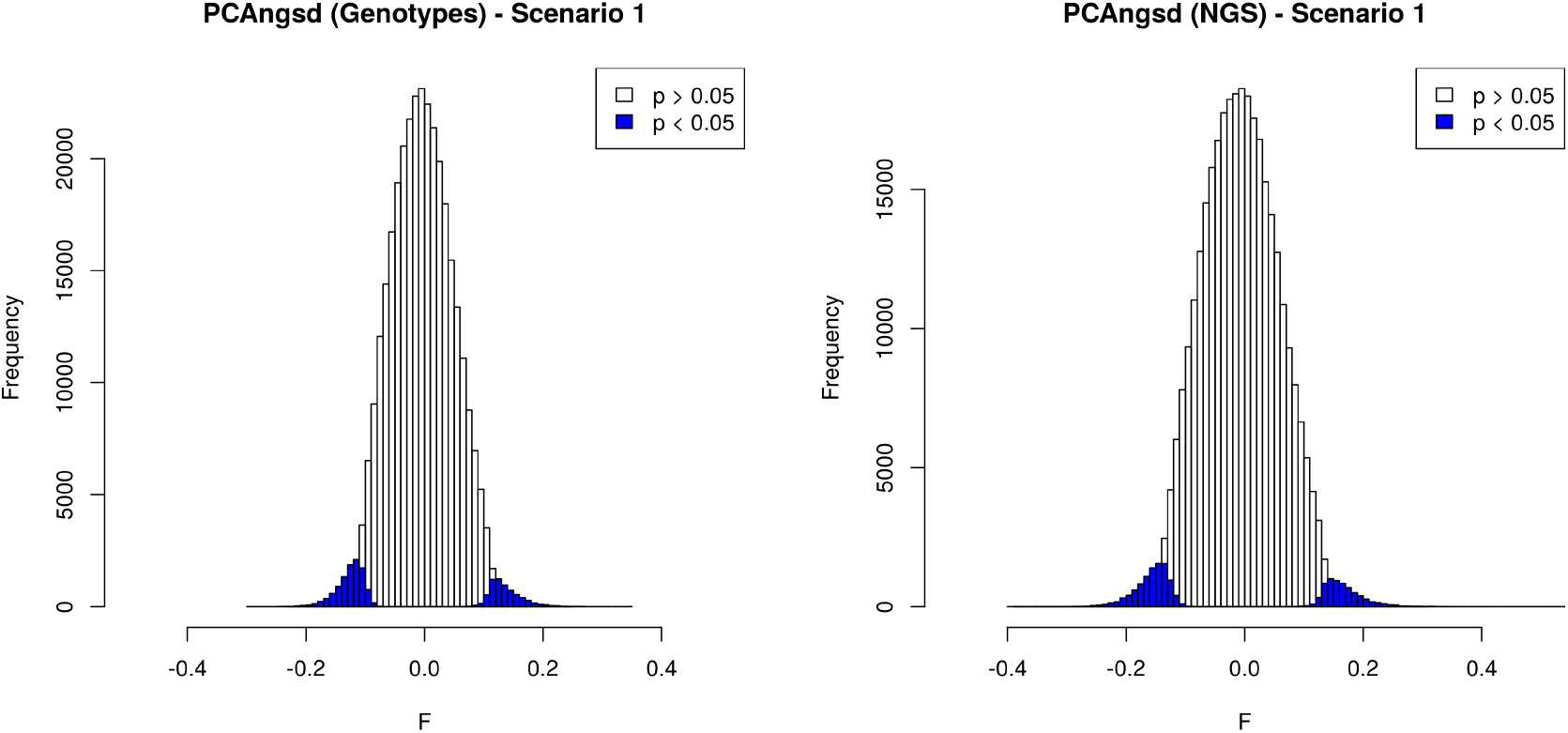
Histograms of the estimated per-site inbreeding coefficients for the simulated scenario with sites sampled from HW proportions. The left plot displays the estimates from simulated genotype data and the right plot shows the estimates from simulated NGS data with a sequencing depth of ∼5X. Sites with a *p*-value lower than 0.05 are colored blue.

**Figure 3:**
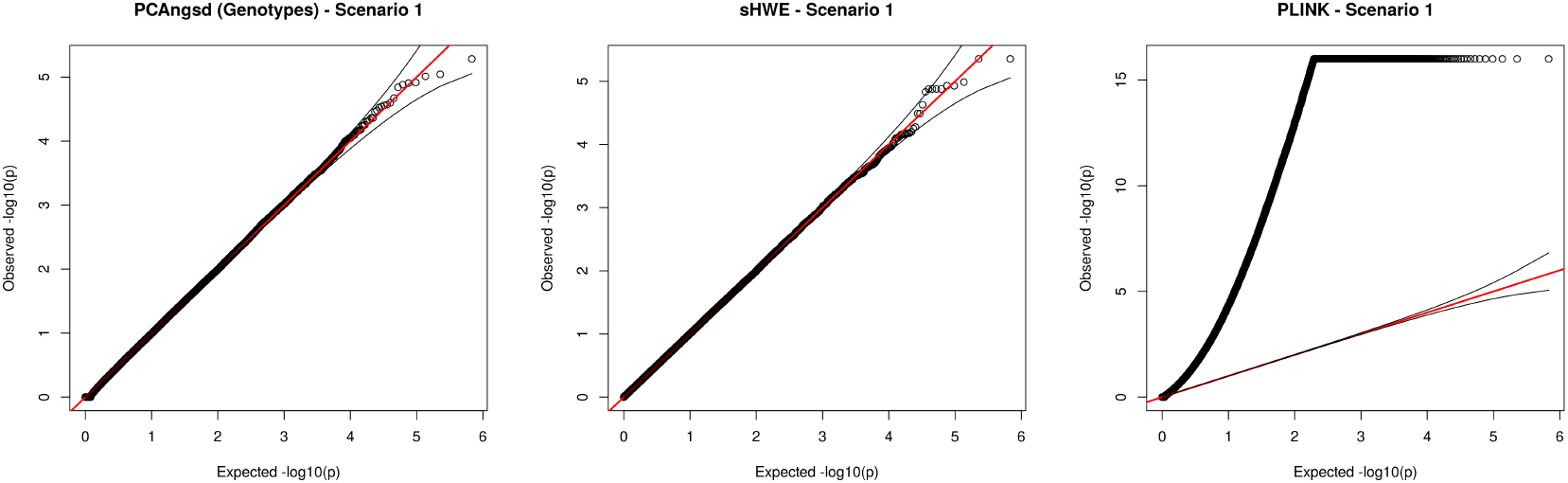
QQ plots of the test statistics in −log_10_ scale using genotype data for Scenario 1, where all sites are sampled from HW proportions. The left plot is the test statistics of PCAngsd, the middle plot of sHWE and the right plot of PLINK.

**Figure 4:**
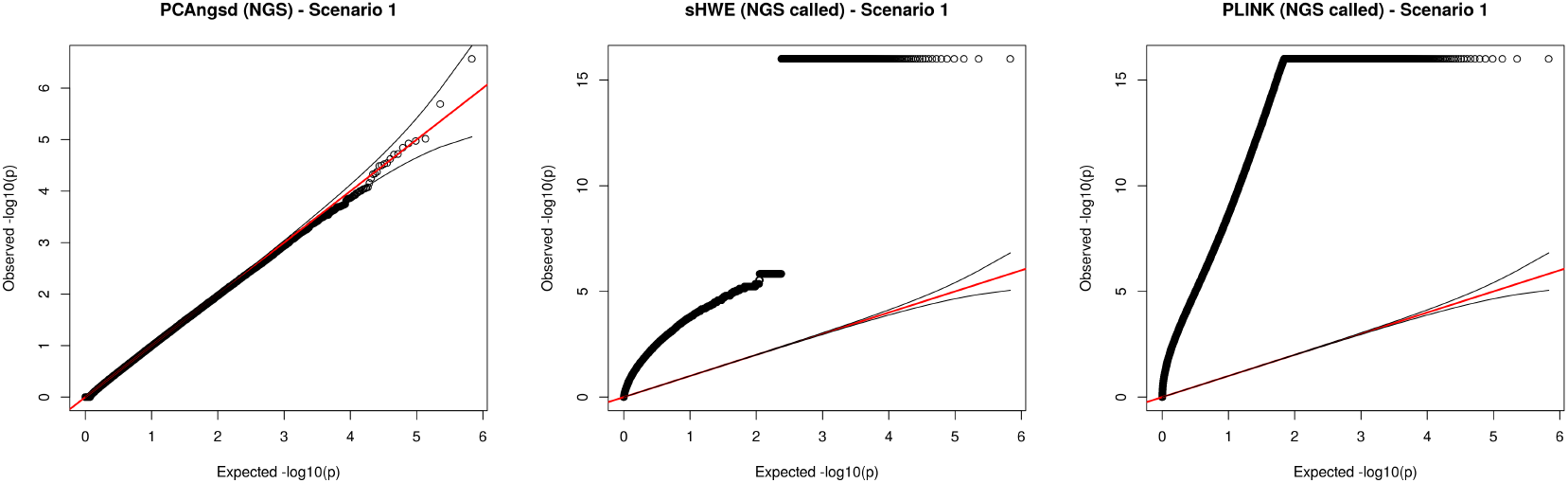
QQ plots of the test statistics in −log_10_ scale using simulated low-depth sequencing data for Scenario 1, where all sites are sampled from HW proportions. The left plot is the test statistics of PCAngsd based directly on genotype likelihoods, while the middle plot of sHWE and the right plot of PLINK are based on naively called genotypes. Due to precision in the outputted *p*-values of sHWE, all *p*-values < 1.0 *×* 10^−16^ are truncated to 16 in −log_10_-scale for convenience in visualization.

When applied to the simulated data in Scenario 2, the effect of negative inbreeding coefficients is seen in the estimates of our method. The estimates for the sites sampled with *F* = −0.5 are slightly biased, as expected, due to the sample frequencies being affected by the negative per-site inbreeding coefficients, as displayed in Figure 5. However, once again the estimates for the sites sampled with *F* = 0 and *F* = 0.5 nicely follow normal distributions around the expected values. Table 1 further show that our method performs well in terms of power and false positive rate (FPR) in comparison to sHWE and PLINK, and it works very well for detecting sites that deviate from HWE with negative per-site inbreeding coefficients. Our method slightly loses power when using low-depth sequencing data but it is still able to keep the expected FPR.

**Figure 5:**
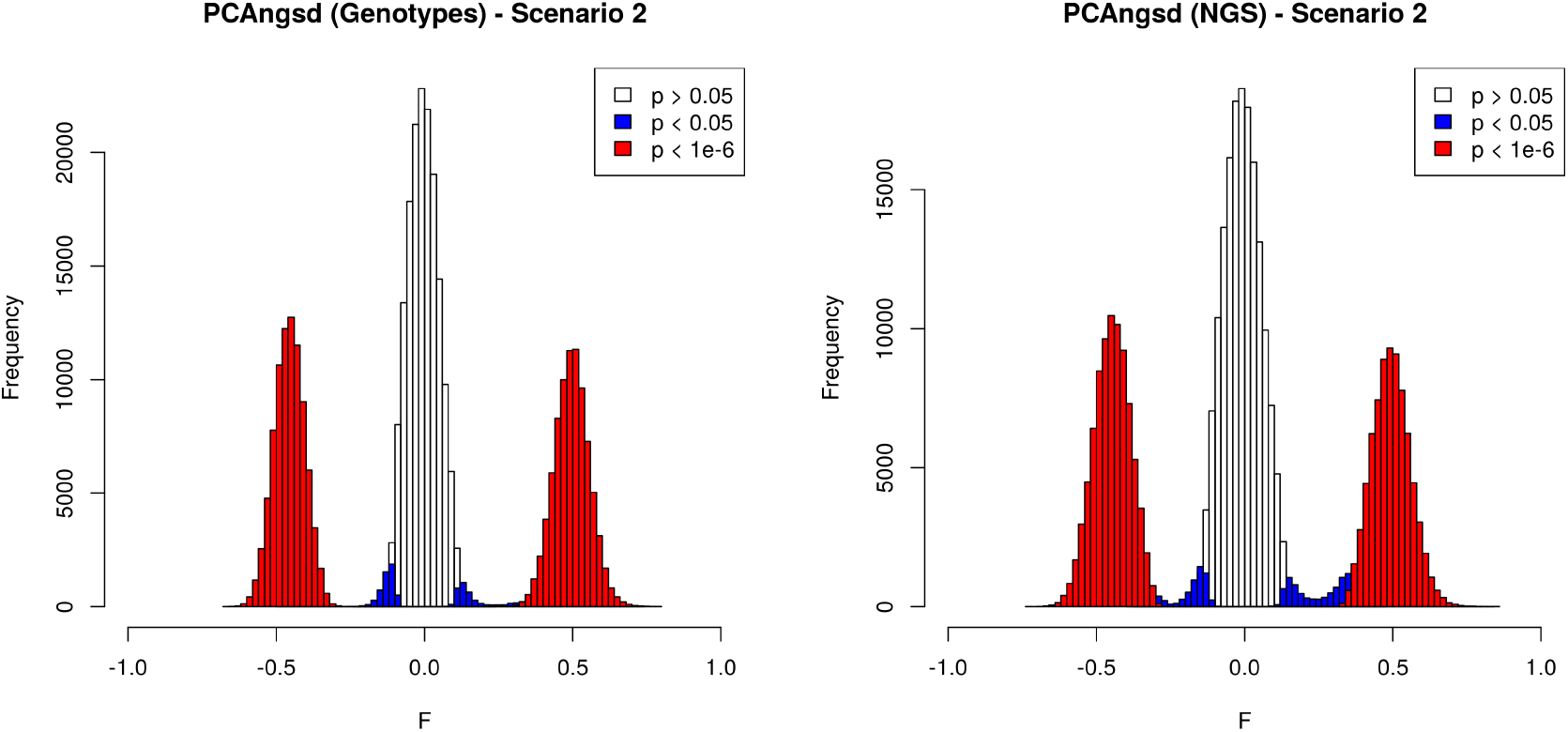
Histograms of the estimated per-site inbreeding coefficients for the simulated scenario with half of the sites sampled with *F* = 0, a quarter sampled with *F* = 0.5 and quarter sampled with *F* = −0.5. The left plot displays the estimates from simulated genotype data and the right plot shows the estimates from simulated NGS data with a sequencing depth of ∼5X. Sites with a *p*-value lower than 0.05 are colored blue, while sites with a *p*-value lower than 1.0 *×* 10^−^6 are colored red.

**Table 1:** Performance of methods on classification of sites out of HW proportions for the simulated dataset with half of its sites sampled with *F* = 0, which are used to measure FPR, a quarter of the sites sampled with *F* = 0.5 and a quarter of the sites sampled with *F* = −0.5, such that half of the sites are out of HW proportions. The two quarters sampled out of HWE are used to measure power. A site is classified as out of HWE for a *p*-value < 1.0 *×* 10^−6^, except for the fourth column where a lower threshold is evaluated. FPR is an abbreviation for the false positive rate.

| Method | Power | FPR | FPR ($\alpha = 0.05$) | Accuracy |
| --- | --- | --- | --- | --- |
| PCAngsd - Genotypes | 0.990 | 0.000 | 0.0511 | 0.995 |
| PCAngsd - NGS | 0.941 | 0.000 | 0.0510 | 0.971 |
| sHWE | 0.988 | 0.000 | 0.0485 | 0.994 |
| PLINK | 0.990 | 0.0589 | 0.575 | 0.965 |

### 3.2 1000 Genomes Project

We also applied our method to genotype and low coverage sequencing data of 366 individuals from four populations in the 1000 Genomes Project (ASW, CEU, CHB, YRI). The two datasets consist of 7.4 million and 7.9 million variable sites, respectively, where 6.9 million of the sites are overlapping. The population structure inferred using PCAngsd is displayed in Figure 6 and once again the top two eigenvectors have been used to model the individual allele frequencies.

**Figure 6:**
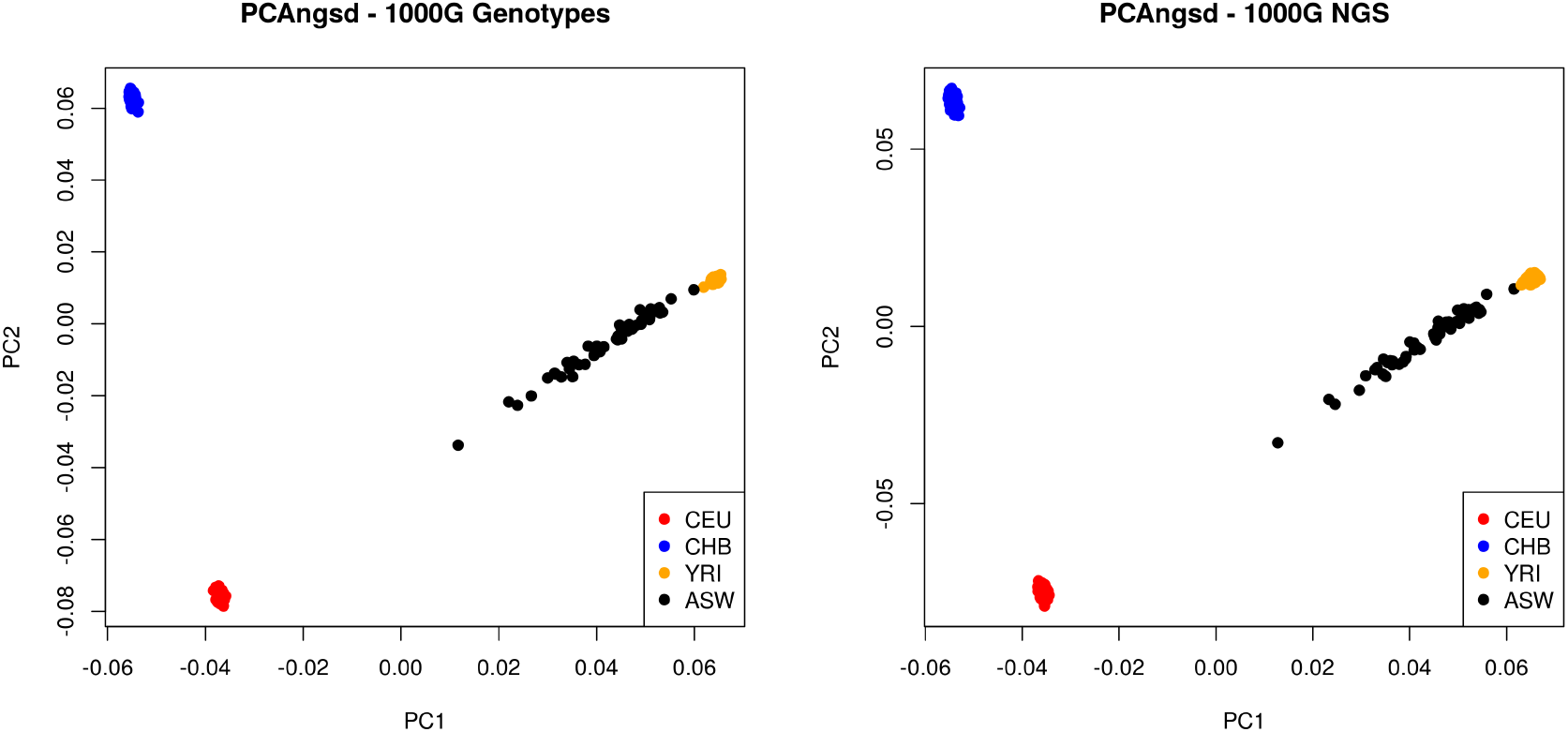
PCA plots of the 1000 Genomes data from 4 different populations. The inferred population structure of the left plot is generated from the genotype dataset, and the right plot is generated from the low coverage sequencing data.

In both datasets, we are able to estimate per-site inbreeding coefficients that follow a normal distribution around 0 (Figure 7), as expected, however the negative estimates are seen to be slightly skewed with a heavy tail for the low coverage sequencing data when using all sites (Figure S2). As seen for the simulations, our test statistic performs very similarly to sHWE using three logistic factors for the genotype data and both methods are able to reduce the number of sites that deviate from HWE by a factor of ∼8 (*α* = 1.0 × 10^−6^) compared to PLINK due to sites assumed to be biased by population structure (Table S1). This effect is considerably smaller for the full set of low coverage sequencing data, where we would also expect to see more technical and random effects causing deviations from HWE than for the genotype data. Though, we show that the performance of PCAngsd, when only analyzing overlapping sites for the the low coverage sequencing data, is very similar to using the genotype data. Most of the significant sites have therefore seemingly been filtered out in the variant calling for the available genotype dataset of the 1000 Genomes Project phase 3 release, thus reinforcing the capabilities of our method in low-depth sequencing data.

**Figure 7:**
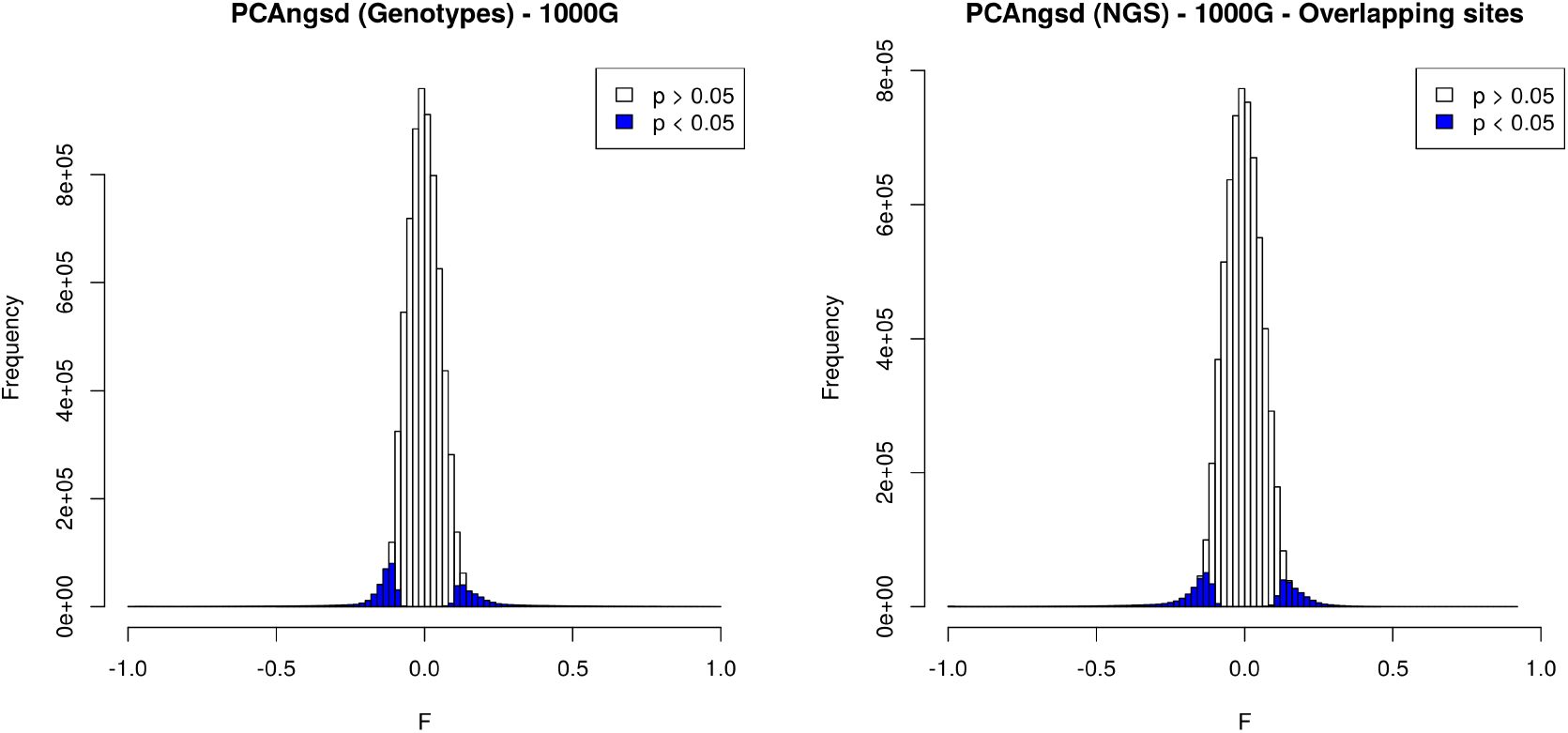
Histograms of the estimated per-site inbreeding coefficients for the 1000 Genomes data from 4 different populations. The left plot displays the estimates from the genotype dataset and the right plot shows the estimates from low coverage sequencing data in overlapping sites. Sites with a *p*-value lower than 0.05 are colored blue.

**Figure 8:**
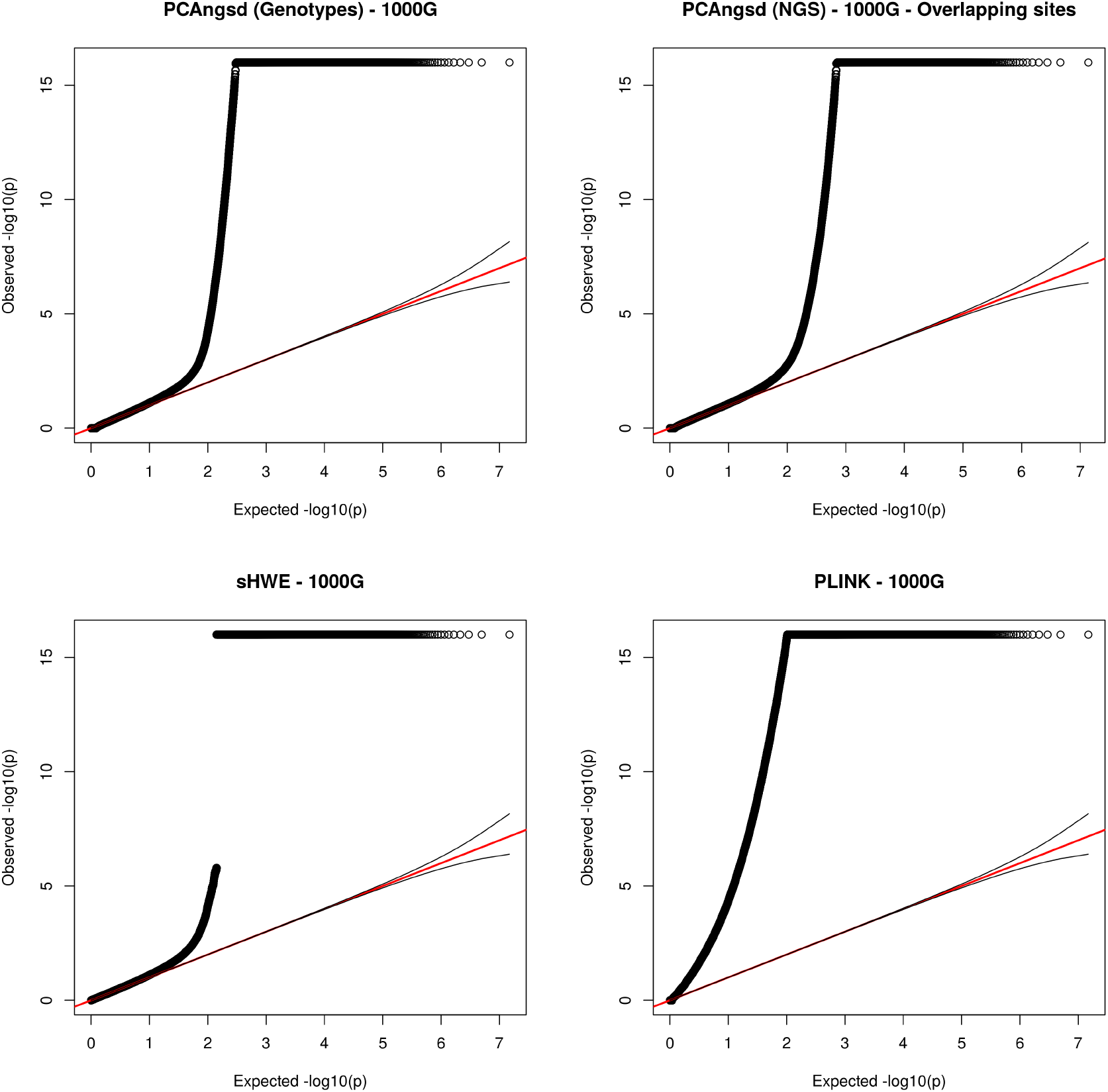
QQ plots of the test statistics in −log_10_ scale for 1000 Genomes Project dataset. The top row displays the plots from PCAngsd based on genotype data for the left plot and low coverage sequencing data for the right plot, using overlapping sites. The bottom row displays the plots from sHWE and PLINK, respectively. Due to precision in the outputted *p*-values of sHWE, all *p*-values < 1.0 × 10^−16^ are truncated to 16 in −log_10_-scale for convenience in visualization.

## 4 Discussion

We have proposed a method to test for HWE in structured populations and integrated it into the PCAngsd framework. This is made possible by incorporating individual allele frequencies, which are modelled from population structure, into a likelihood framework that estimate deviations from HWE. The method is able to work on both sequencing data, in the form of genotype likelihoods in Beagle format, and genotype data in binary PLINK format.

We have applied our method to both simulated and real data to test for HWE in structured populations, where we demonstrate that our method is performing well using both sequencing and genotype data in the presence of population structure. The simulation studies show that our method performs very similarly to an existing method for genotype data, sHWE, where both methods are able to detect deviations from HWE caused by other factors than population structure. However, our method also performs well on simulated low-depth sequencing data (∼5X) where we are able to keep the statistical power high while keeping the false-positive rate low. The bias of calling genotypes for low-depth sequencing data has also been demonstrated using sHWE and PLINK, but PCAngsd is able to overcome this bias by working directly on genotype likelihoods.

Our presented results for the 1000 Genomes Project datasets are not as clean as seen in the study of Hao and Storey [24], as we analyze all variable sites across the whole genome without filtering for sites overlapping the genotyping chip from the phase 3 release. We further show that a lot of the sites that deviates from HWE in the full low coverage sequencing data are not present in the phase 3 variant callset of the 1000 Genomes Project. Thus verifying the usefulness of PCAngsd, as it is able to detect the deviations from HWE due to technical errors when using low-depth sequencing data in structured populations.

In addition to testing for deviations from HWE, PCAngsd estimates per-site inbreeding coefficients that quantify the deviation from HWE and are useful for understanding why sites are out of HWE. We have proposed a heuristic extension to a likelihood framework such that we are able to estimate negative per-site inbreeding coefficients. In this way, we can model deviations from HWE in both directions that may provide deeper insight into sites of evolutionary interest regardless of observing excess of homozygosity or excess of heterozygosity in structured populations.

## 5 Funding

This project was funded by the Lundbeck foundation (R215-2015-4174).

